## Supplement for "The effect of acute oral galactose administration on the redox system of the rat small intestine"

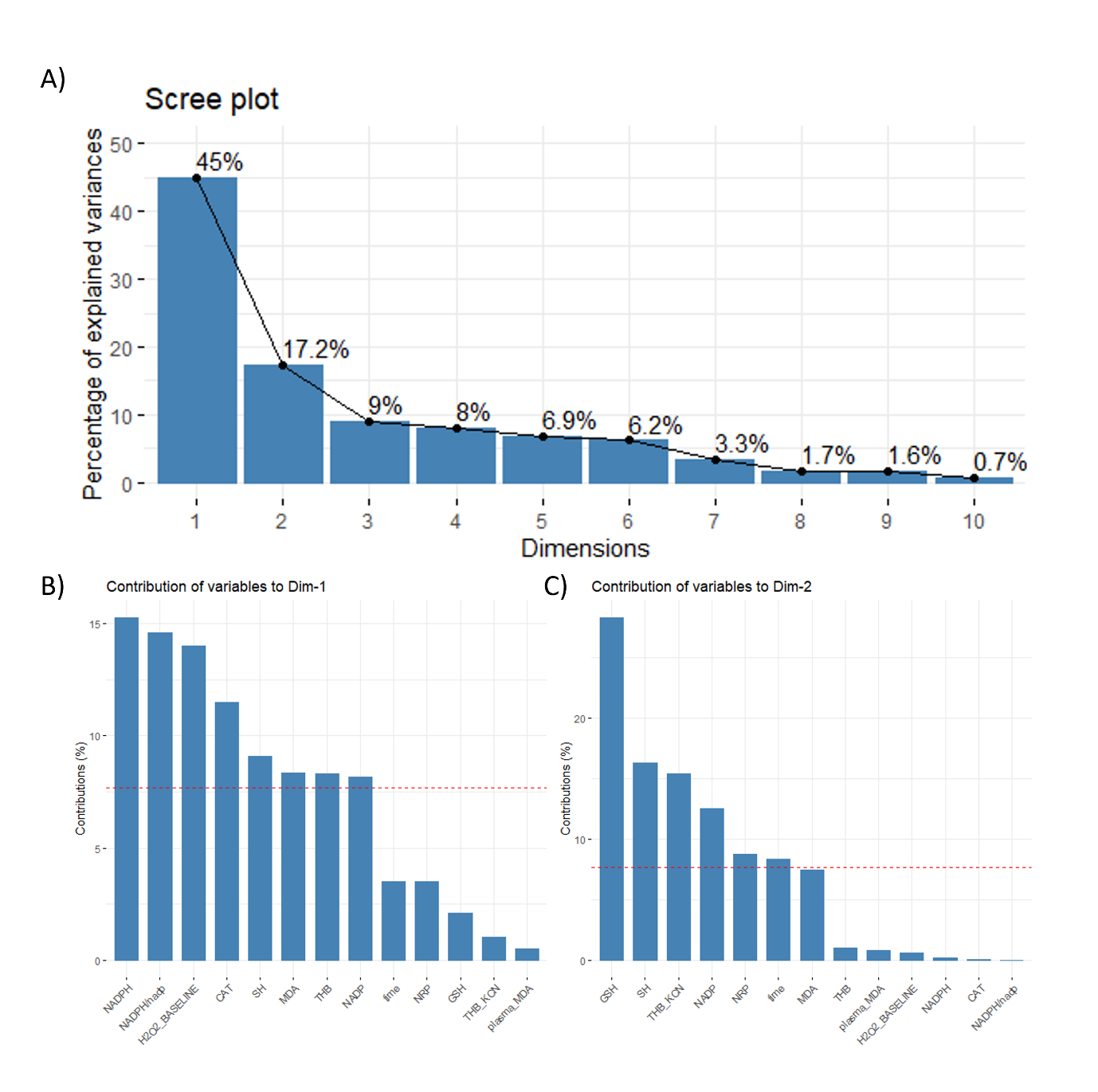

**Fig S1.** Combined intestinal redox-related parameters principal component analysis. A) Scree plot. B) Contribution of individual variables to the 1^st^ principal component. C) Contribution of individual variables to the 2^nd^ principal component. NRP – nitrocellulose redox permanganoemtry; time – minutes since the exposure to treatment (continuous variable); THB_KCN – change in 1,2,3-trihydroxybenzene autooxidation rate in the presence of 2mM potassium cyanide (KCN); GSH – glutathione (the main representative of the low molecular weight thiols (LMWT)); MDA – malondialdehyde (the main representative of the thiobarbituric acid reactive substances (TBARS)); NADP – nicotinamide adenine dinucleotide phosphate; THB – change in 1,2,3-trihydroxybenzene autooxidation rate; CAT – catalase activity (hydrogen peroxide dissociation rate); H2O2_BASELINE – estimated baseline levels of tissue hydrogen peroxide; SH – protein reactive sulfhydryl groups; NADPH – reduced nicotinamide adenine dinucleotide phosphate; NADPH/nadp – the concentration of reduced nicotinamide adenine dinucleotide phosphate corrected for concentration of nicotinamide adenine dinucleotide phosphate; contrib – contribution of variables; Dim1 – 1^st^ dimension; Dim2 – 2^nd^ dimension.

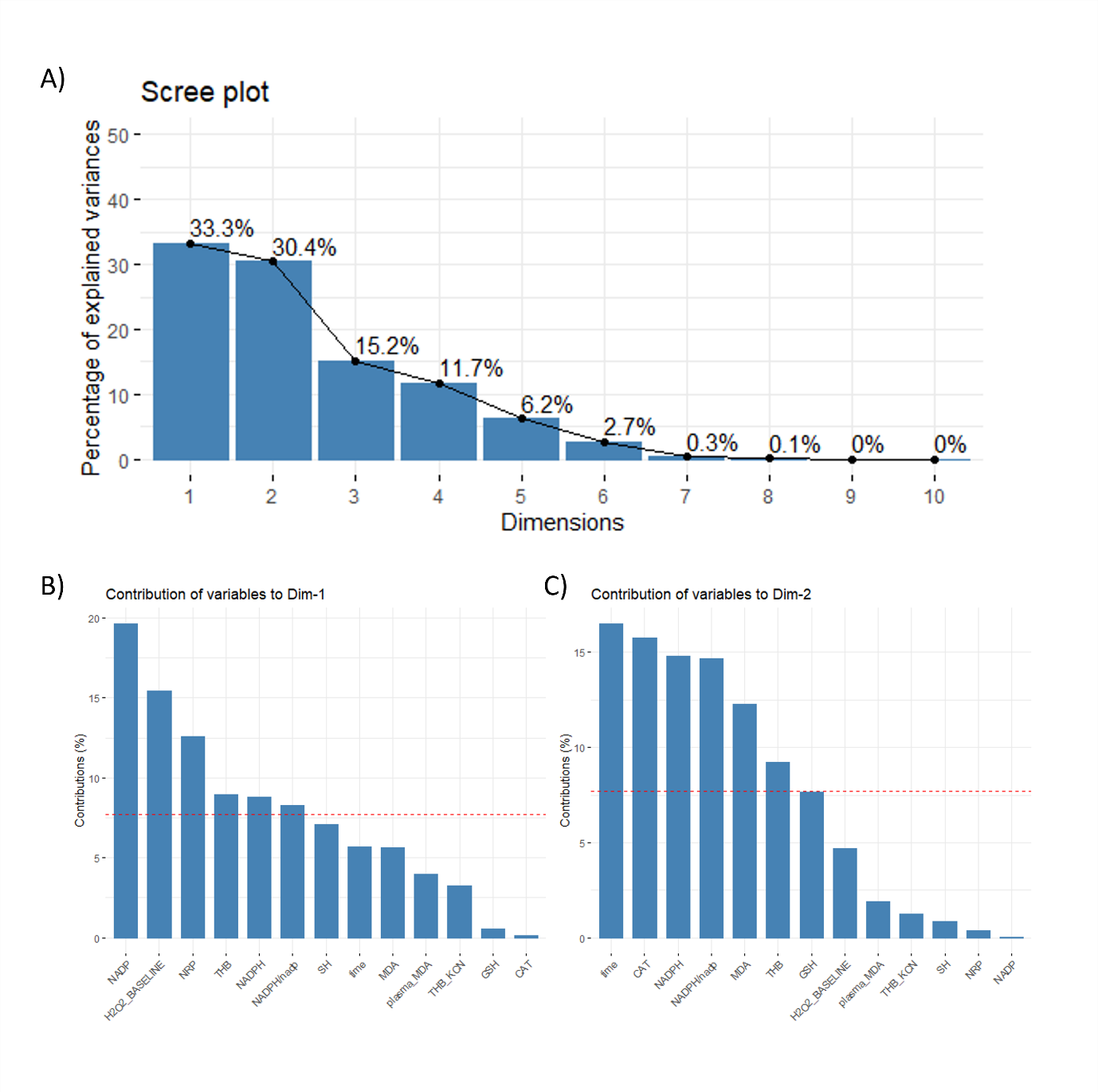

**Fig S2.** Duodenal redox-related parameters principal component analysis. A) Scree plot. B) Contribution of individual variables to the 1^st^ principal component. C) Contribution of individual variables to the 2^nd^ principal component. NRP – nitrocellulose redox permanganoemtry; time – minutes since the exposure to treatment (continuous variable); THB_KCN – change in 1,2,3-trihydroxybenzene autooxidation rate in the presence of 2mM potassium cyanide (KCN); GSH – glutathione (the main representative of the low molecular weight thiols (LMWT)); MDA – malondialdehyde (the main representative of the thiobarbituric acid reactive substances (TBARS)); NADP – nicotinamide adenine dinucleotide phosphate; THB – change in 1,2,3-trihydroxybenzene autooxidation rate; CAT – catalase activity (hydrogen peroxide dissociation rate); H2O2_BASELINE – estimated baseline levels of tissue hydrogen peroxide; SH – protein reactive sulfhydryl groups; NADPH – reduced nicotinamide adenine dinucleotide phosphate; NADPH/nadp – the concentration of reduced nicotinamide adenine dinucleotide phosphate corrected for concentration of nicotinamide adenine dinucleotide phosphate; contrib – contribution of variables; Dim1 – 1^st^ dimension; Dim2 – 2^nd^ dimension.

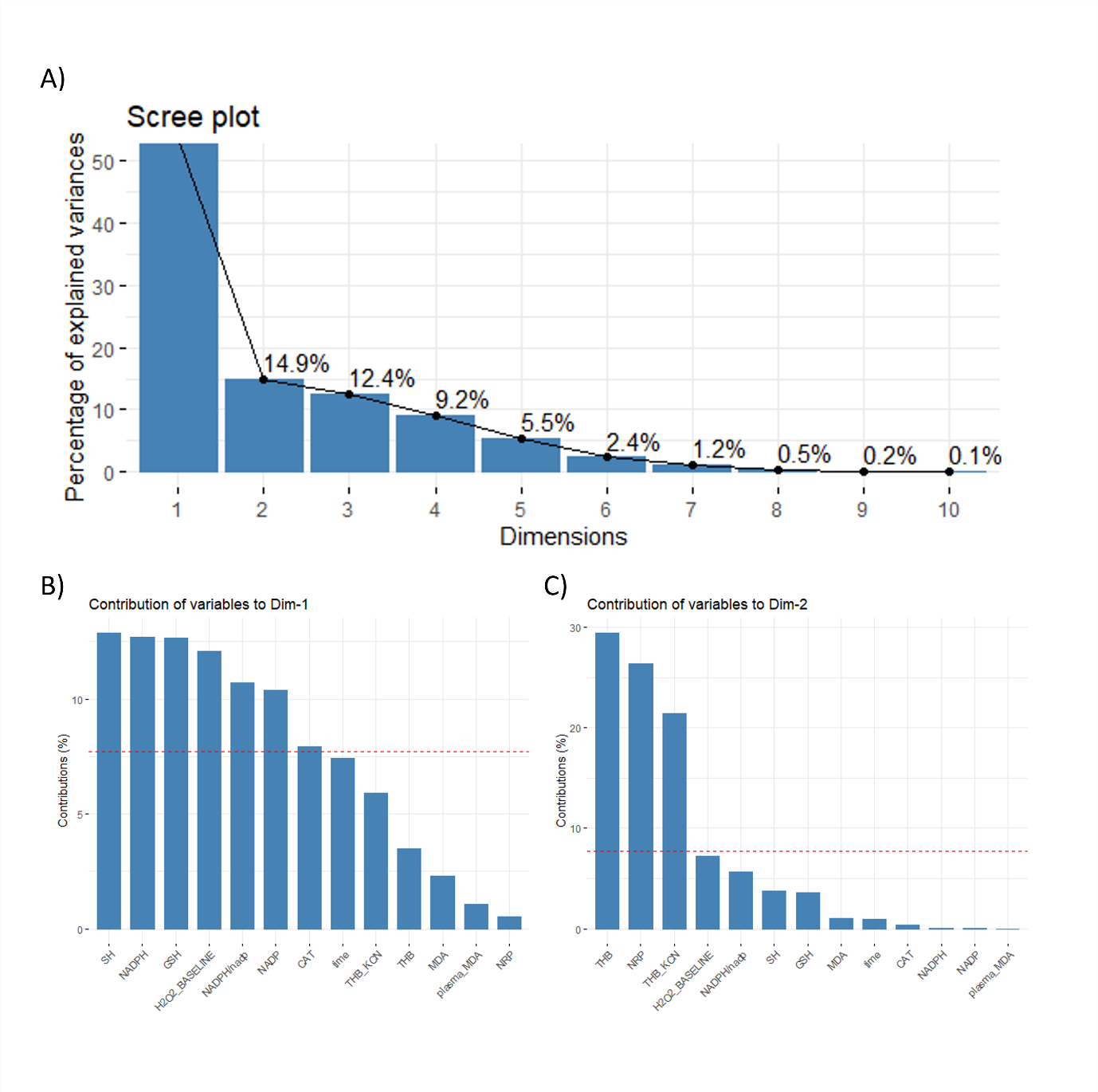

**Fig S3.** Ileal redox-related parameters principal component analysis. A) Scree plot. B) Contribution of individual variables to the 1^st^ principal component. C) Contribution of individual variables to the 2^nd^ principal component. NRP – nitrocellulose redox permanganoemtry; time – minutes since the exposure to treatment (continuous variable); THB_KCN – change in 1,2,3-trihydroxybenzene autooxidation rate in the presence of 2mM potassium cyanide (KCN); GSH – glutathione (the main representative of the low molecular weight thiols (LMWT)); MDA – malondialdehyde (the main representative of the thiobarbituric acid reactive substances (TBARS)); NADP – nicotinamide adenine dinucleotide phosphate; THB – change in 1,2,3-trihydroxybenzene autooxidation rate; CAT – catalase activity (hydrogen peroxide dissociation rate); H2O2_BASELINE – estimated baseline levels of tissue hydrogen peroxide; SH – protein reactive sulfhydryl groups; NADPH – reduced nicotinamide adenine dinucleotide phosphate; NADPH/nadp – the concentration of reduced nicotinamide adenine dinucleotide phosphate corrected for concentration of nicotinamide adenine dinucleotide phosphate; contrib – contribution of variables; Dim1 – 1^st^ dimension; Dim2 – 2^nd^ dimension.
